# MolUNet++: Adaptive-grained Explicit Substructure and Interaction Aware Molecular Representation Learning

**DOI:** 10.1101/2025.07.07.663425

**Authors:** Fanding Xu, Zhiwe Yang, Wu Su, Lizhuo Wang, Deyu Meng, Jiangang Long

## Abstract

Molecular representation learning is a critical task in AI-driven drug development. While graph neural networks (GNNs) have demonstrated strong performance and gained widespread adoption in this field, efficiently extracting and explicitly analyzing functional groups remains a challenge. To address this issue, we propose MolUNet++, a novel model that employs Molecular Edge Shrinkage Pooling (MESPool) for hierarchical substructure extraction, utilizes a Nested UNet framework for multi-granularity feature integration, and incorporates a substructure masking explainer for quantitative fragment analysis. We evaluated MolUNet++ on tasks including molecular property prediction, drug-drug interaction (DDI) prediction, and drug-target interaction (DTI) prediction. Experimental results demonstrate that MolUNet++ not only outperforms traditional GNN models in predictive performance but also exhibits explicit, intuitive, and chemically logical interpretability. This capability provides valuable insights and tools for researchers in drug design and optimization.

## Introduction

Molecular representation serves as a crucial link connecting various scientific disciplines to machine learning, including bioinformatics, cheminformatics, materials science, and particularly drug discovery (1–3). Accurate and task-related representation is essential for predicting molecular properties, behaviors, and interactions (4). Although traditional molecular representations, such as handcrafted molecular fingerprints and descriptors, have been widely utilized in drug discovery, they often face limitations in terms of generalizability, expressiveness, and the complexity of the molecular structures they can effectively capture (5–7).

Molecular representation learning (MRL) is a specialized field that utilizes deep learning techniques to extract task-relevant features directly from raw input data, such as SMILES strings and molecular graphs. This capability enables the discovery of intricate patterns that may remain elusive to traditional descriptors (8). A significant advancement in MRL has been the emergence of Graph Neural Networks (GNNs), as molecules can naturally be represented as graphs.

GNNs excel at learning both local and global structural features through the iterative aggregation of information from neighboring nodes (atoms) and edges (bonds) (9, 10). GNN-based methods have demonstrated superior performance in various molecular tasks, including property prediction, interaction prediction, and dynamics simulations (11–13).

However, despite addressing certain limitations of traditional methods by automatically capturing topological and relational information within molecular graphs, GNNs share the same interpretability challenges as other deep learning models (14). The predictions generated by GNNs often lack a clear correlation to specific molecular features, which impedes a comprehensive understanding of how structural elements affect molecular properties.

In molecular science, the relationship between a molecule’s structure and its properties is a fundamental principle. This principle asserts that the specific arrangement of atoms within a molecule, including aspects such as functional groups and bond types, significantly influences its physical and chemical behaviors (15). However, most message passing based GNNs obscure the structural reasoning behind their predictions, making it challenging for scientists to understand the connection between a molecule’s structure and its predicted properties.

Graph pooling has emerged as a powerful method for extracting hierarchical substructural features from molecular graphs at various scales. By progressively reducing the size of the graph while preserving essential substructural information, graph pooling has been shown to enhance models’ ability to capture substructures relevant to specific tasks (16–18). Despite its significant potential to identify topological substructures, most graph pooling models still struggle to provide explicit, interpretable representations that align with domain-specific intuition. This often results in the loss of important fine-grained structural information, yielding only coarsegrained representations at the final stage (19).

To address these challenges, we introduce MolUNet++ (Figure 1), a novel GNN-based MRL architecture. Building on our previous work, which introduced an attention-based graph pooling method, MESPool (20) (Figure 1c), which explicitly identifies task-related substructures in molecules, thereby enhancing interpretability and performance compared to traditional pooling methods. We have designed a corresponding unpooling module (Figure 1d), utilizing a GNN architecture inspired by Nested UNet (21) to connect hierarchical node features (Figure 1a). This results in adaptive representations at the atomic level and facilitates the construction of a pretraining pipeline tailored for property prediction. Additionally, we have incorporated an interactionguided attention mechanism into the pooling process to improve substructure identification, thereby enhancing MRL for tasks such as drug-drug interaction (DDI) and drug-target interaction (DTI) predictions. We further deployed a structural masking explainer to quantitatively analyze the molecular fragments identified by MolUNet++ (Figure 1e). In our experiments, we compared MolUNet++ with state-of-theart (SOTA) GNN-based methods, demonstrating that MolUNet++ not only improves predictive performance but also exhibits interpretability consistent with principles of structural chemistry. To further evaluate its effectiveness, we selected two representative DDI and DTI models that employ GNNs as molecular encoders. By replacing the original GNN encoders with MolUNet++, we conducted performance analyses. The results indicated that MolUNet++ not only achieved superior performance but also provided more chemically intuitive molecular explanations compared to the original attention-based interpretive mechanisms. These findings highlight the enhanced structural awareness of MolUNet++ over traditional GNNs, offering valuable insights for researchers in compound optimization and drug design.

**Fig. 1.**
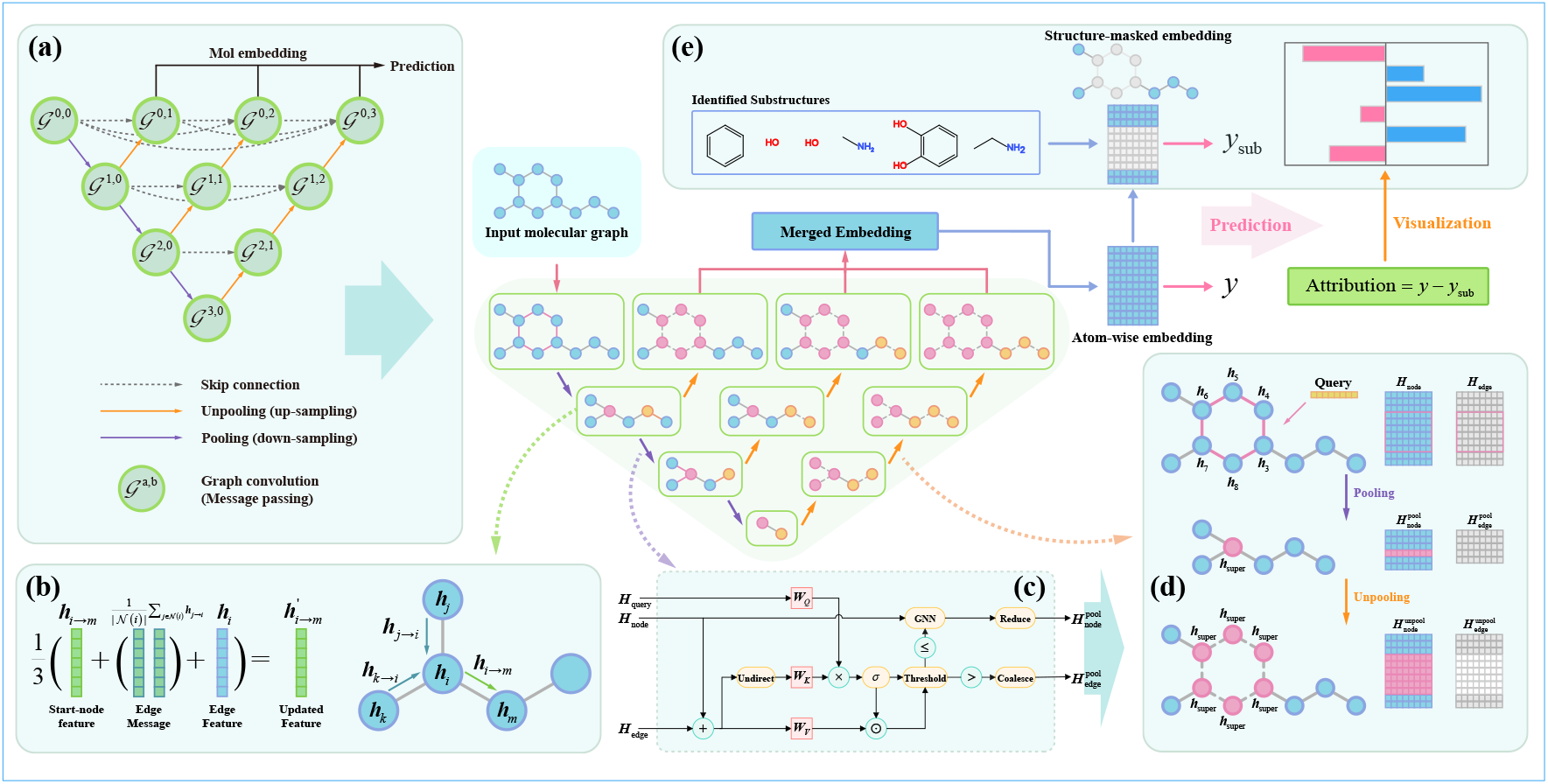
Overview of the MolUNet++ architecture. (a) The architecture of MolUNet++ incorporates pooling and unpooling operations as down-sampling and up-sampling processes, respectively. Each layer following the unpooling operation retains feature representations from the previous layers of the same scale through skip connections. Ultimately, molecular embeddings at various granularities are combined for prediction tasks. (b) The edge feature update process within the edge message passing (E-MP) module. (c) The pipeline of the MESPool process: node features and edge features are integrated to represent undirected bonds, which are then scored by a query feature. The edges are classified as inner-substructure and backbone bonds, with nodes clustered accordingly and shrunk into super-nodes. (d) An illustration of the pooling and unpooling operations applied to the dopamine toy structure. The unpooling operation reconstructs the original graph structure, copies and aligns the super-node feature to its member nodes, and resets the features of the inner-substructural edges to zero vectors. (e) The substructure explainer: MolUNet++ encodes atom features and merges them into a molecular representation through global pooling, subsequently predicting the property *y*. By masking the atom features of substructures identified by MESPool, a new prediction, *y*_sub_, is obtained for the molecule with those substructures removed. The contribution of each substructure to the property is determined by *y* − *y*_sub_, and this contribution is visualized to provide insights.

## Methods

### MESPool

MESPool is designed under the Selection, Reduction and Connection (SRC) pooling scheme (18). In the selection phase, an edge scoring function is employed to assign weights to edges. Low-scored and high-scored edges are distinguished using a threshold function, categorizing them as inner-substructure bonds and backbone bonds, respectively. During the reduction phase, nodes connected by low-scored edges are shrunk into super-nodes, effectively reducing the graph’s complexity while preserving essential structural features.

Since MESPool requires scoring dynamical edge features, while most MP layers do not include edge feature updates, here we defined an edge message passing (E-MP) operator 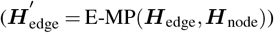 before pooling to update edge features and integrate them with node features (Figure 1b):

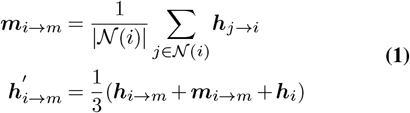

where ***h***_*i→m*_ and ***m***_*i→m*_ denotes the feature and message of the directed edge *i → m* respectively.

### Pooling

Taking the node features ***H***_node_ ∈ ℝ ^*N ×d*^ and the edge features ***H***_edge_ ∈ ℝ^*M ×d*^ as input, ***H***_edge_ is firstly updated via E-MP. Here we introduce an additional query feature 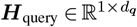 to flexibilize the structure identification in MESPool (Figure 1d), which is defined by the mean feature of all nodes (Equation 2) as default:

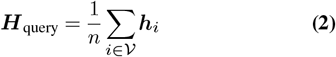

Note that ***H***_query_ can be also defined as the feature of another entities in various tasks, such as the feature of another compound in DDI tasks, or the protein feature in DTI tasks (see Figure 2). The edge attention can then be calculated following the functions:

**Fig. 2.**
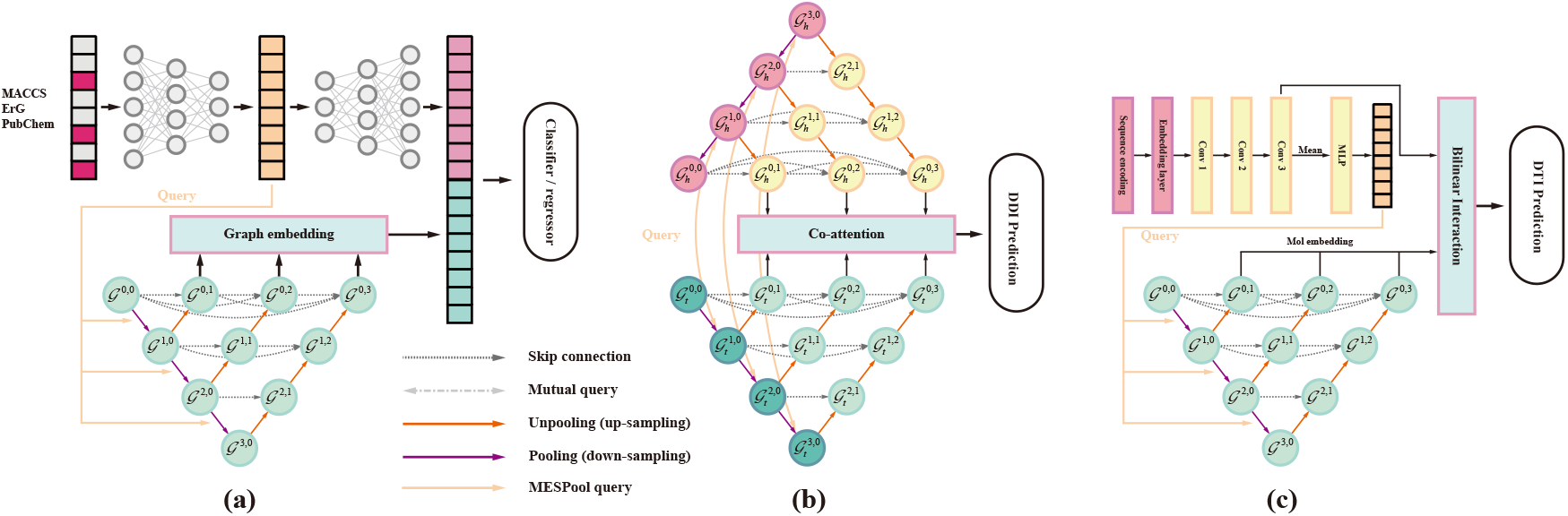
Utilization of structurual identification query (Equation 3) across different tasks. (a) Molecular property prediction: Three molecular structural fingerprints are concatenated and encoded using a multi-layer perceptron (MLP). The resulting feature serves as the query feature **H**_query_ in MESPool, and is subsequently decoded by another MLP for property prediction in conjunction with the graph embedding. (b) A pair of molecules is encoded using the same MolUNet++ network, with each molecule mutually utilizing the other’s graph features as **H**_query_. The output embeddings are integrated through a co-attention layer to predict DDI interactions. (c) DTI prediction: The protein sequence is encoded using 1D CNNs, and the average of the representation rows is transformed through an MLP to serve as the **H**_query_ for structural identification in the drug encoder. The drug and protein representations are then fed into a bilinear attention network to learn their pairwise local interactions.

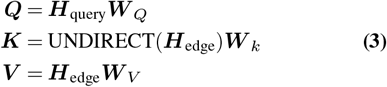

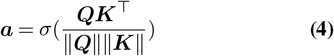

where 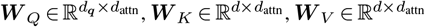 are learnable weight matrices, and ***a*** ∈ ℝ^*M*^ is the calculated edge attention scores. The UNDIRECT(*•*) converts the edge features to undirected by averaging the features of two opposing edges, and *σ*(•) denotes the sigmoid activation function. The edge features are subsequently updated using the attention scores to facilitate gradient propagation: ***H***_edge_ = ***a*** · ***K***.

A dual-controlled threshold is used to distinguish edges into two categories, which is defined as the minimum value of a manually set hyperparameter *λ* and the mean value of ***a***: threshold = MIN(*λ*, MEAN(***a***)). We consider the low-scored edges and connected nodes to be the innersubstractural components, and group those into subsets with a strongly connected component finding algorithm (22):

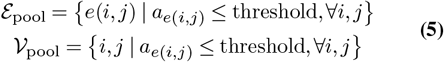

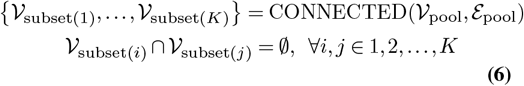

where *a*_*e*(*i,j*)_ is an element in ***a***. An additional GNN layer is adopted on the subsets to learn the representation of the substructures, and the subsets can be further shrunk into supernodes with the updated node features:

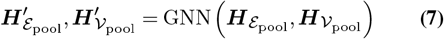

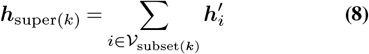

Some nodes may be connected by multiple backbone edges after the shrunk, those redundant edges can be coalesced the with a simple summation function, and eventually a reduced graph is obtained.

### Unpooling

We propose an unpooling operator of MESPool as the up-sampling process for pooled molecular graphs. The unpooling process is relatively straightforward, as illustrated in Figure 1b. It begins by restoring the graph to its pre-pooled connectivity structure. Specifically, each super-node, which represents a pooled substructure, is decomposed back into its original member nodes. The features of the super-node are then copied and assigned to each of its constituent members, ensuring consistency of the node-level features within the restored graph (Equation 9).

Furthermore, the edge features within the super-nodes (i.e., the inner-substructural edges) are reset to zero vectors (see Equation 10). Subsequently, another E-MP operator is applied to update the global edge features (Equation 1). Zeroing the inner-substructural edge features erases the substructure-specific edge information and eliminates their contribution to the backbone edges in the E-MP. Conversely, these inner-substructural edges are initialized with new features derived from the messages propagated through the backbone edges.

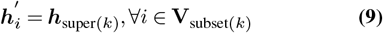

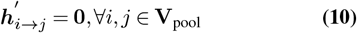

### MolUNet++

UNet++ (21), an extension of UNet (23), was originally developed for medical image segmentation. It refines the encoder-decoder architecture through multi-stage down-sampling and up-sampling, thereby minimizing information loss during the down-sampling process. Furthermore, its dense connectivity mechanism facilitates more effective feature propagation across different levels of the network, improving feature integration and segmentation accuracy. GraphUNet (24), which is widely utilized for graph-related tasks, employs TopK pooling (25) for down-sampling, focusing on sparse nodes but often overlooks their topological relationships. Although effective for creating hierarchical graph representations, this approach restricts the preservation of detailed local information, rendering it less suitable for tasks such as molecular representation learning.

The architecture of MolUNet++ is illustrated in Figure 1a. In this framework, we utilize MESPool and its unpooling operators as multi-stage processes for down-sampling and up-sampling, which correspond to multi-grained substructures. Node-level features of varying granularity, yet within the same graph scale, are interconnected through nested skip pathways. The skip connection is applied following the unpooling process and is defined as:

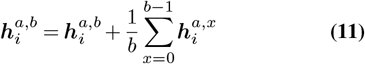

where 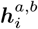 represents the feature of node *i* in the graph at scale *a* and granularity *b*. This nested skip mechanism allows the model to effectively integrate multi-grained features, ultimately yielding an adaptive-grained representation of the graph.

MolUNet++ generates node-level and edge-level graph representations corresponding to the number of pooling layers *L*. For graph-level molecular prediction tasks, it is crucial for performance to effectively merge these multi-grained node features and integrate them into graph-level representations. We refer to these two operations as Jumping Knowledge (JK) and Global Pooling (GP):

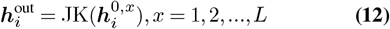

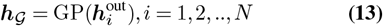

where *L* represents the number of pooling layers (the depth of the model), ***h***_*𝒢*_ denotes the graph embedding. Additionally, the selection of GNN layers within the model is also significant. Ultimately, we selected concatenation for JK, summation for GP, and PNA (26) as the graph convolution layer in the model. A detailed discussion on operator selection and the corresponding ablation study is provided in the Supporting Information.

### Pretrain strategy for molecular prediction prediction

Graph pretraining for molecular tasks has gained traction as a method to enhance the representation of molecular graphs. Recent studies have demonstrated that combining self-supervised learning (SSL) and supervised learning (SL) strategies can yield superior pretraining performance compared to using either approach in isolation (27–30). Building on this concept, we designed a two-stage pretraining process: Stage 1 employs SSL on the **ZINC 15** dataset (200 million unlabeled compounds) (27, 31) to optimize both node-level and edge-level representation capabilities. In Stage 2, we utilize SL to incorporate domain-specific knowledge using the **ChEMBL** dataset (over 456k compounds with 1310 types of bioassay binary labels) (32, 33).

Stage 1 and Stage 2 are performed sequentially. After pretraining, the model parameters are fine-tuned for downstream property prediction tasks using a reduced learning rate to preserve the features acquired during pretraining.

#### Stage 1

Attribute Masking (AttrMask) is a pioneering approach in graph self-supervised learning (SSL) (27). A typical AttrMask method involves randomly masking nodes or edges and training the model to predict the masked node or edge types using their encoded embeddings. In this study, we adopt two improved masking strategies: weighted attribute masking (34) and motif masking (35), focusing on atom-level and motif-level structural information respectively. In addition, we introduce an auxiliary SL classification task to predict molecular decomposition information. Inspired by the SME method (14), we label each node/atom in the molecule according to functional groups, BRICS decomposing (36), and Murcko scaffolds(37). For functional groups, we take the 39 important groups from **RDKit** and use them to match each atom.

We adopt cross-entropy (CE) and binary cross-entropy (BCE) loss function for the SSL and SL task, respectively (see Supporting Information for the detail of the loss functions), the total loss for stage 1 can be described as:

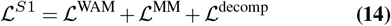

where ℒ^WAM^, ℒ^MM^ and ℒ^decomp^ denotes the loss for weighted attribute masking, motif-aware masking and auxiliary decomposition classification tasks, respectively.

#### Stage 2

In stage 2, we focus on optimizing graph-level features by introducing domain knowledge through SL. Specifically, we use the labeled ChEMBL dataset to train the model to predict binary labels for 1,310 experimental bioassays. We also pre-computed 200 physical and chemical descriptors for each molecule (38), and the model is trained to predict these descriptors through regression.

Similar to stage 1, here we also introduce a auxiliary SSL task in stage 2. For a batch of molecules, we calculate the cosine similarity of their graph embeddings and fit it to the Tanimoto similarity of their RDKit molecular fingerprints. We use ℒ^sim^, ℒ^exp^, and ℒ^cal^ represent the loss of the molecular similarity, bioassay classification and descriptor regression (see Supporting Information for the detail of the loss functions), the total loss of the stage 2 can be then described as:

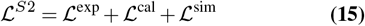

### Substructure explainer

The atom representations encoded by MolUNet++ are used to predict molecular properties through a decoding process (GP followed by a classifier / regressor), resulting in the prediction of the property *y*. We construct a set of molecular fragments from all the substructures identified by the pooling layers. Masking the atom representations of the fragment and re-decoding the molecular representation by:

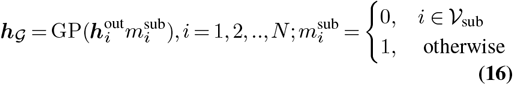

we obtain a new prediction *y*_sub_ for the molecule with the fragment removed. We define an attribution value as:

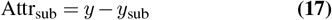

which reflects the contribution of a specific substructure to the molecular property. The sign of the attribution value indicates whether the substructure tends to increase or decrease the overall property value, while its magnitude represents the strength of the effect.

## Results

In this study, we evaluate the performance of the MolUNet++ architecture in molecular property prediction tasks, comparing it with traditional GNN models. A two-stage pretraining strategy is designed for MolUNet++ to incorporate relevant domain-specific knowledge. Furthermore, we replace the traditional GNN molecule encoder with MolUNet++ in drug-drug interaction (DDI) and drug-target interaction (DTI) tasks with two well-known models, evaluating its performance in these contexts. In addition, we investigate the structural cognitive interpretability of the query-guided pooling mechanism within MolUNet++, incorporate structural molecular fingerprints, drug, and protein features, to guide the identification of structures and interactions across these tasks (Figure 2). All experiments were conducted on a hardware platform comprising an AMD 5950X CPU, an RTX 4090D GPU, and 64GB of RAM. The software platform was developed using **RDKit**^1^, **PyTorch**^2^, and **PyG**^3^.

### Molecular property prediction

The property prediction experiments utilized 8 representative datasets from **MoleculeNet**^4^, which include 5 classification datasets: **BBBP, BACE, Clintox, Tox21**, and **Sider**, as well as 3 regression datasets: **Lipophilicity, FreeSolv**, and **ESOL**. For evaluation metrics, we employed the Area Under the Receiver Operating Characteristic Curve (AUC-ROC) for classification tasks and the Root Mean Squared Error (RMSE) for regression tasks. To establish a comprehensive benchmark, we compared MolUNet++ against a basic GNN model, GraphUNet (24), and several well-established pretrained models, including AttrMask (27), ContextPred (27), GraphMAE (39), Mole-BERT (40), GraphCL (28), and MGSSL (41), utilizing the hyperparameter settings as specified in their original publications. For each dataset and model, we deploy the same scaffold splitting (37), and take 5 independent runs to calculate the average performance and standard deviation as the results. Detailed information regarding hyperparameter selection for our experiments is available in the Supporting Information.

### Benchmark

The performance results for property prediction are summarized in Table 1 and Figure 3a. Among the non-pretrained models, GraphUNet outperformed the Basic GNN model across multiple tasks, while MolUNet++ demonstrated the highest performance across six datasets, particularly in BACE, Lipophilicity, and ESOL. Furthermore, MolUNet++ exhibited significantly stronger performance than the Basic GNN model in seven tasks, excluding only BBBP, and even outperformed most pretrained models in certain tasks. Notably, the pretrained MolUNet++ (MolUNet++ Pretrain) showed performance improvements across all tasks, highlighting the effectiveness of pretraining in enhancing model generalizability.

**Table 1.**
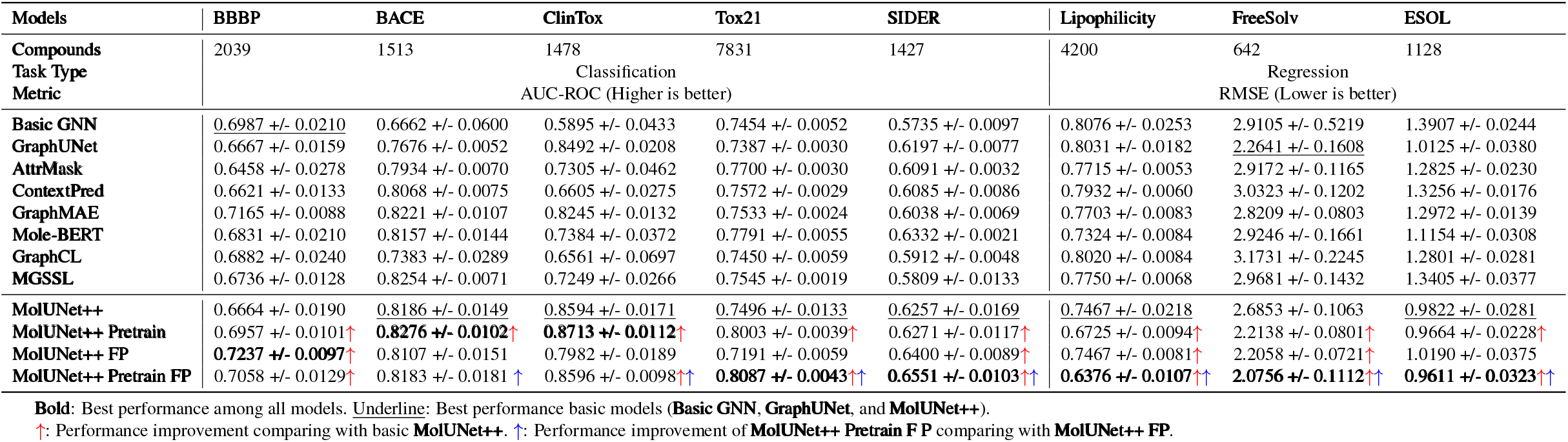
Benchmark performance on molecular property prediction datasets.

**Fig. 3.**
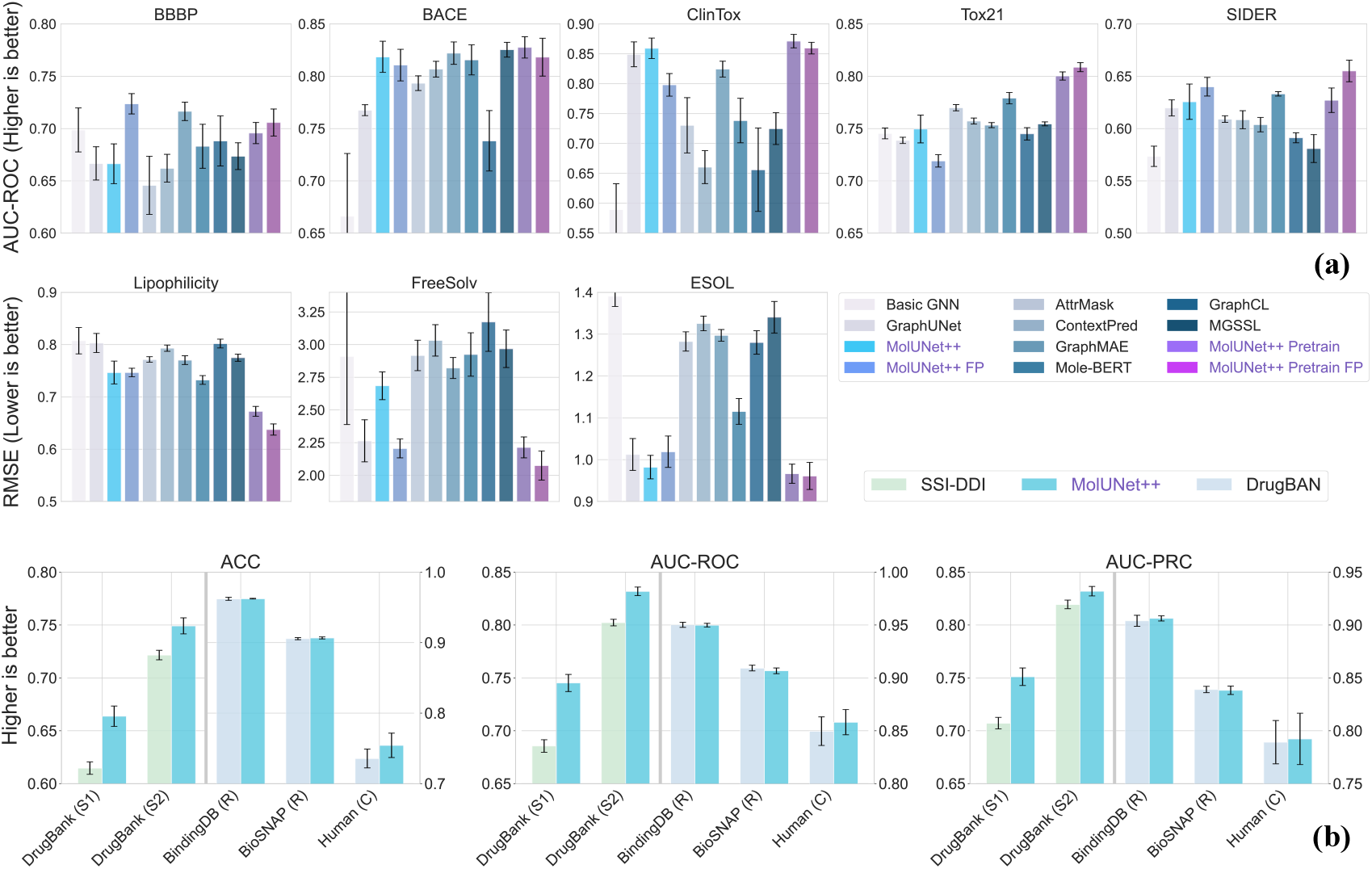
Benchmark performance. (a) Property prediction: the figures compare non-pretrained models (the first 4 bars) with pretrained models (the last 8 bars). The first 5 charts utilize AUC-ROC as the metric, where <u>higher bars indicate better performance</u>. The last 3 charts employ RMSE, where <u>lower bars indicate better performance</u>. (b) Interaction prediction: in each figure, the left two bar groups show DDI performance, while the right three show DTI performance. Each figure represents a different metric, where <u>higher bars indicate better performance</u>

To demonstrate the capability of MESPool in leveraging external features for guiding structural identification, we also explored the use of molecular structural fingerprints as the query feature in MESPool (MolUNet++ FP and MolUNet++ Pretrain FP). The framework of the model utilizing fingerprints is illustrated in Figure 2a, with additional details provided in the Supporting Information. Although the incorporation of molecular fingerprints did not result in significant performance improvements across all datasets, it did enhance prediction accuracy on certain datasets, particularly BBBP. Furthermore, we observed a consistent trend with the original models, where the pretrained versions outperformed their non-pretrained counterparts. Ultimately, the pretrained MolUNet++ model with molecular fingerprints achieved the highest performance across most datasets. Overall, MolUNet++ consistently surpassed the baseline models on all eight datasets.

### Interpretability

In MolUNet++, MESPool explicitly identifies substructures, and the output atom representations incorporate information about these substructures. To quantify the contribution of these substructures to property predictions, we employ an explanation mechanism similar to the Substructure Masking Explainer (SME) (14) (see Figure 1e). We construct a set of molecular fragments from all the substructures identified by the pooling layers and calculate their contributions to the predictions (see Equation 17).

In this section, we present several molecular examples to demonstrate the interpretability of MolUNet++ on the benchmark dataset Lipophilicity (distribution coefficient, log *D*), and the Acute Toxicity LD50 dataset (42), which includes 7,385 compounds, with the 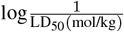 value serving as the label. The higher the dose, the more lethal of a drug. We utilize the best-performing model configuration to conduct the following analyses (the training results for the LD50 dataset are provided in the Supporting Information). The results are illustrated in Figure 4, the attribution values in the figures are normalized to range between 0 and 1.

**Fig. 4.**
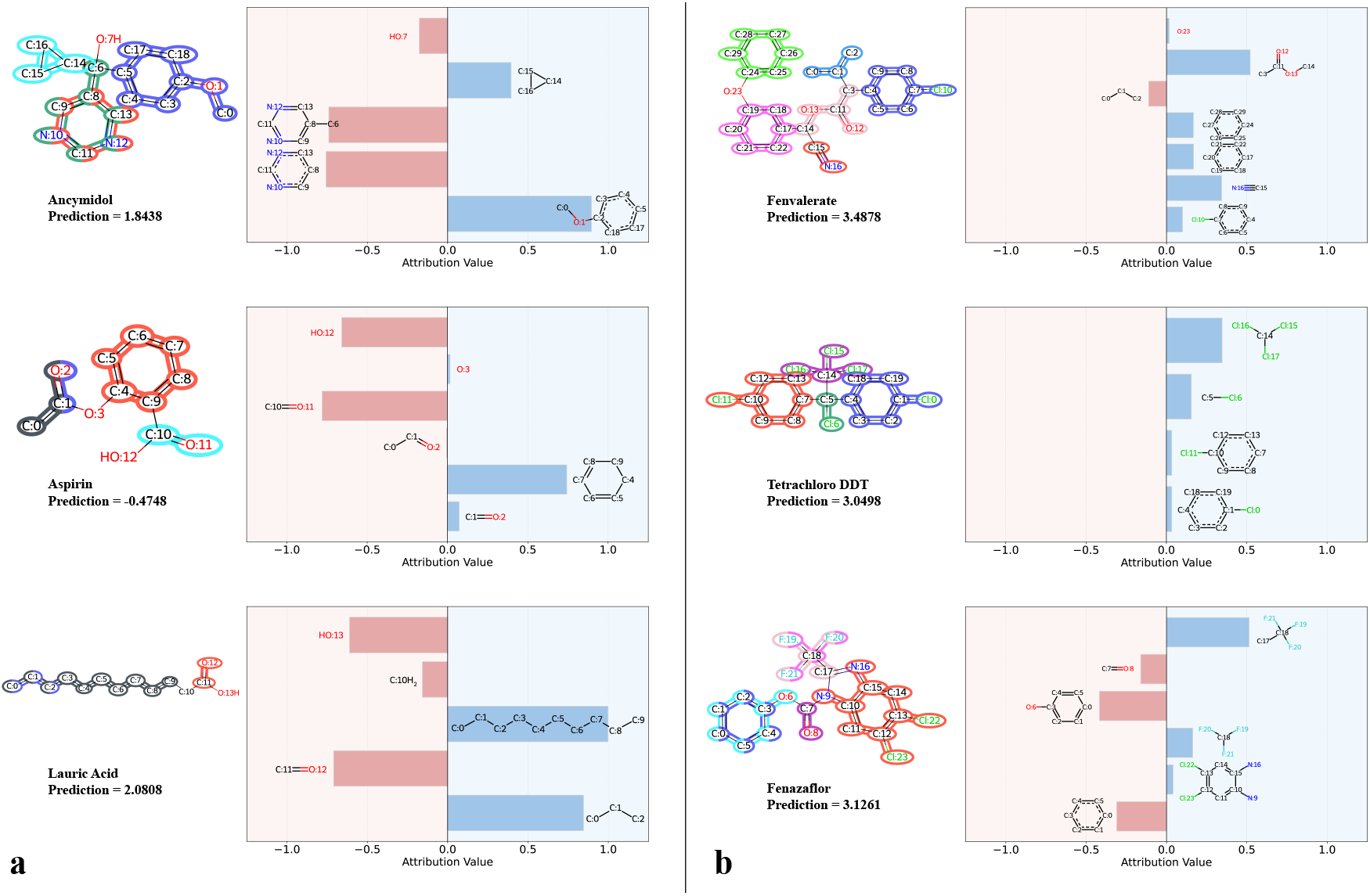
Visualization of pooling results and substructure attribution values in 2 property prediction tasks. Atoms of the same color represent an identified fragment, while independent atoms are also regarded as a single atomic substructure. (a) log *D* prediction: positive values indicate an increase in hydrophobicity (lipophilicity), whereas negative values indicate the contrary; (b) LD_50_ toxicity prediction: positive values suggest increased lethality.

The interpretability of MolUNet++ in predicting the distribution coefficient (log *D*) is consistent with the study of substituent contributions to log *D* conducted by Landry et al. (43). For example, the model identifies the negative contributions of the pyrimidine ring and hydroxyl group, as well as the positive contribution of the cyclopropane group in Ancymidol, which align well with the reported Δlog *D* in their study. Similarly, for Lauric Acid, MolUNet++ recognizes the positive contribution of the long carbon chain to hydrophobicity, which consequently diminishes the polar contribution of the carboxyl. Interestingly, in the case of Aspirin, the model identifies distinctly different attribution values for the C=O structures present in the carboxyl and ester groups. This distinction corresponds to the differing polarities of these groups, with the C=O in the carboxyl group exhibiting greater hydrophilicity compared to that in the ester group. This finding indicates that MolUNet++ effectively utilizes hierarchical information within the molecular structure, capturing subtle differences in group contributions.

In the context of toxicity prediction, MolUNet++ effectively identifies classical toxicophores, including cyano groups, chlorine substituents (such as the tetrachloroethyl bridge in Tetrachloro DDT), and fluorine substituents (such as the trifluoromethyl group in Fenazaflor). These well-known toxic groups are assigned relatively high attribution values. Furthermore, we observe significant differences in the contributions of the benzene rings in Fenvalerate and Fenazaflor. This discrepancy arises because Fenvalerate serves as a pyrethroid ester insecticide and acaricide, with phenoxybenzyl and carboxylic ester groups being typical structures of pyrethroids. As a result, the two benzene rings in Fenvalerate exhibit high attribution values. This observation highlights the capability of MolUNet++’s pooling and unpooling mechanism to capture information at varying structural granularities.

### Interaction prediction

To further emphasize the superior molecular encoding capabilities of MolUNet++ compared to general GNNs, we conducted experiments on both DDI and DTI tasks. Specifically, we selected two well-established models for these tasks: **SSI-DDI** (44) for DDI prediction and **DrugBAN** (45) for DTI prediction. Both models utilize traditional standalone GNNs as molecular encoders, enabling us to directly compare the performance of MolUNet++ with that of GNNs by substituting the encoder in these frameworks while keeping all other components unchanged. In these tasks, unlike property prediction, we utilize the features of interacting molecules and proteins as queries within the MESPool mechanism. To ensure a fair comparison of performance between MolUNet++ and traditional GNNs for accessing their efficacy for interaction prediction, we strictly adhered to the parameter settings reported in the original studies for SSI-DDI and DrugBAN. The performance results are presented in Table 2 and Figure 3.b

**Table 2.** Performance comparison of the original SSI-DDI and DrugBAN and encoder replaced version in DDI and DTI tasks.

| Interaction | Dataset | Models | ACC | AUC-ROC | AUC-PRC |
| --- | --- | --- | --- | --- | --- |
| DDI | DrugBank (S1) | SSI-DDI | 0.6147 +/- 0.0059 | 0.6856 +/- 0.0059 | 0.7072 +/- 0.0055 |
|  |  | MolUNet++ | **0.6638 +/- 0.0096** | **0.7452 +/- 0.0081** | **0.7511 +/- 0.0083** |
|  | DrugBank (S2) | SSI-DDI | 0.7216 +/- 0.0045 | 0.8023 +/- 0.0031 | 0.8195 +/- 0.0041 |
|  |  | MolUNet++ | **0.7492 +/- 0.0076** | **0.8319 +/- 0.0040** | **0.8320 +/- 0.0045** |
| DTI | BindingDB (R) | DrugBAN | 0.9622 +/- 0.0021 | 0.9502 +/- 0.0024 | 0.9040 +/- 0.0052 |
|  |  | MolUNet++ | 0.9624 +/- 0.0005 | 0.9499 +/- 0.0017 | 0.9063 +/- 0.0025 |
|  | BioSNAP (R) | DrugBAN | 0.9057 +/- 0.0015 | 0.9093 +/- 0.0027 | 0.8390 +/- 0.0030 |
|  |  | MolUNet++ | 0.9067 +/- 0.0014 | 0.9067 +/- 0.0027 | 0.8383 +/- 0.0040 |
|  | Human (C) | DrugBAN | 0.7357 +/- 0.0134 | 0.8496 +/- 0.0135 | 0.7892 +/- 0.0205 |
|  |  | MolUNet++ | **0.7543 +/- 0.0173** | **0.8581 +/- 0.0119** | **0.7923 +/- 0.0243** |

(S1): Cold-start split S1 in DrugBank dataset. (S2): Cold-start split S2 in DrugBank dataset. (R): Random split in DTI dataset. (C): Cold-start split in DTI dataset.

#### DDI

SSI-DDI employs a GAT network to independently encode two molecules, with each layer outputting a graphlevel feature 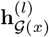 and 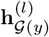, which aligns with the multigranularity graph-level representations of MolUNet++. The features are weighted using a Co-attention layer and further combined to make the interaction prediction with a RESCAL model (46).

We replaced GAT with MolUNet++ (Figure 2b), using the same network depth and hidden feature dimension while keeping the predictor unchanged. For benchmarking, we used the widely recognized DDI dataset **DrugBank**^5^, applying the two cold-start splits S1 and S2 mentioned in the original study:

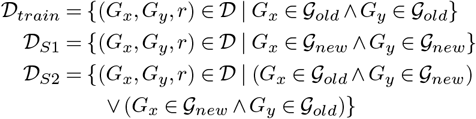

_𝒢 *new*_ denotes the set of new drugs (unseen during training), and 𝒢 _*old*_ denotes the seen drugs, such that 𝒢 _*new*_ ∪ 𝒢 _*old*_ = 𝒢 and 𝒢 _*new*_ ∩ 𝒢 _*old*_ = ∅. Both the original SSI-DDI and the modified version is trained on _*train*_ and tested on both 𝒟_*S*1_ and 𝒟_*S*2_. The results are shown in Table 2, it can be observed that, under both splits, MolUNet++ achieved significant performance improvements compared to the original SSI-DDI.

In the interpretability case study, we followed the examples provided in SSI-DDI: Phosphodiesterase-5 (PDE5) inhibitors, which are commonly used to treat erectile dysfunction, are contraindicated for co-administration with organic nitrate drugs due to the risk of a synergistic reduction in blood pressure. For visual analysis, we selected the molecular representations with the highest pairwise interaction scores from the co-attention layer. Specifically, for SSI-DDI, we visualized the atom attention scores derived from the selected molecular representations of the corresponding SAGPooling (47) layer in the GAT encoder, where higher scores are indicated by darker colors. For MolUet++, we applied the same approach as in property prediction: we masked the substructures identified by the corresponding MESPool layer in the selected molecular representations to calculate attribution values. Fig. 5 illustrates the visualization results, highlighting that, compared to the node attention mechanism in the original SSI-DDI, MolUNet++ demonstrates a more chemically intuitive understanding of substructures. It accurately and comprehensively identifies all nitrate groups while showcasing their positive contributions to the interaction.

**Fig. 5.**
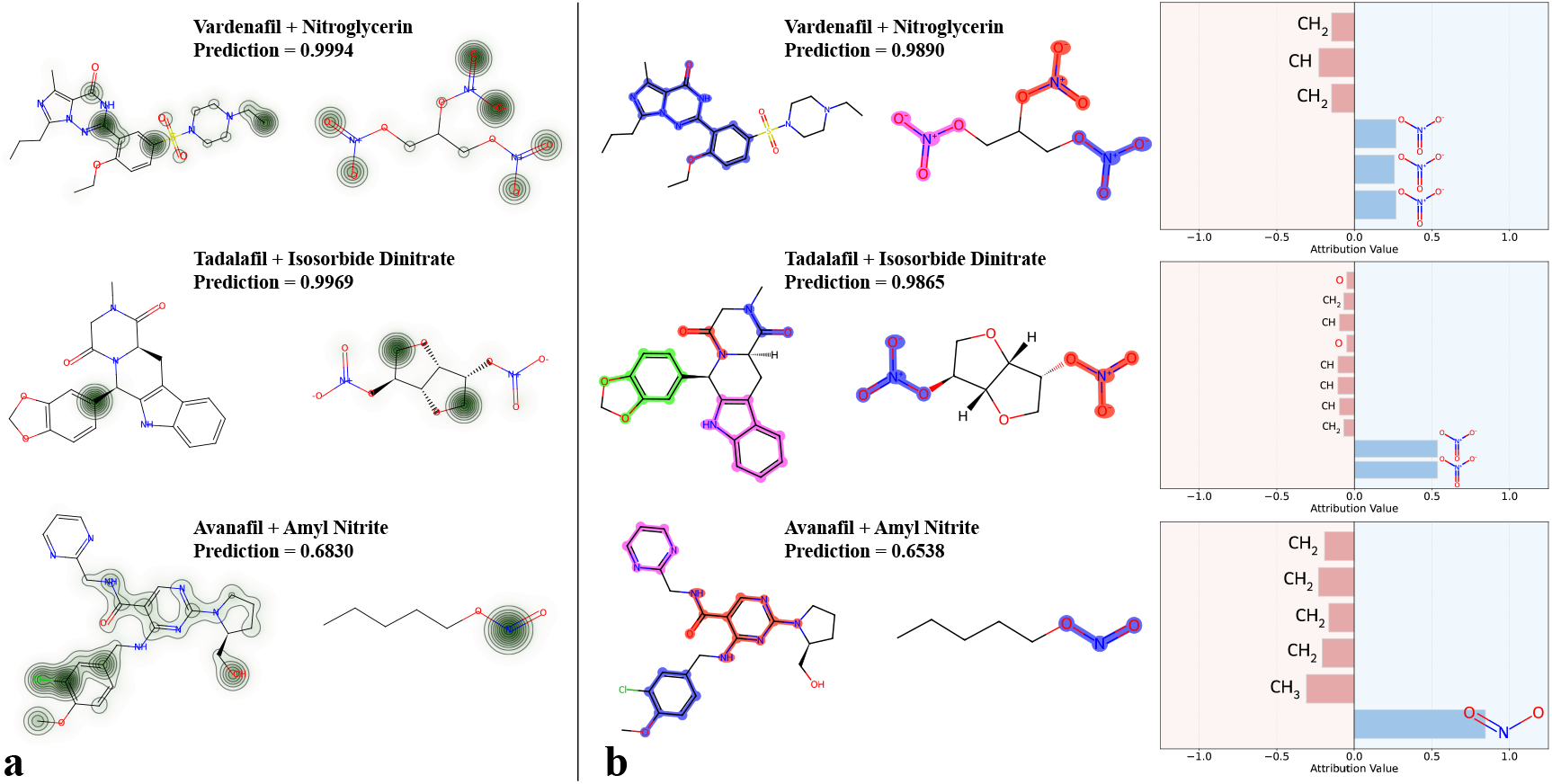
Visualization of the case study in DDI prediction. (a) Attention scores visualization in the original SSI-DDI; (b) pooling results and identified substructure contribution visualization in MolUNet++.

### DTI

Given an input drug–target pair, DrugBAN utilizes separate Graph Convolutional Network (GCN) and onedimensional Convolutional Neural Network (1D CNN) blocks to encode molecular graph and protein sequence information, respectively. A bilinear attention network is employed to encode the interaction, generating a joint representation for DTI prediction. We replaced the GCN encoder with our MolUNet++, while keeping the other parts of the model unchanged. The modified model framework is shown in Figure 2c.

We conducted benchmark tests on two randomly split datasets, BindingDB (48) and BioSNAP (49), as well as a cold-start dataset, Human (50) (where neither the proteins nor the drugs in the test set appeared in the training set). On the two randomly split datasets, the performance of MolUNet++ was comparable to that of the original model. We attribute this similarity to the lack of challenge in the random splits, where the results approached the performance ceiling. However, in the cold-start task, MolUNet++ demonstrated a noticeable performance improvement. This indicates that MolUNet++, with its enhanced structural awareness, achieves better generalization capabilities compared to the GCN in scenarios involving novel drugs.

To analyze molecular-level interpretability, we followed the case study presented in DrugBAN and conducted a visualization analysis using two co-crystallized ligands from the PDB database: 6QL2 (ethoxzolamide complexed with human carbonic anhydrase II) and 5W8L (9YA ligand bound to human L-lactate dehydrogenase A). DrugBAN highlights the top 20% weighted atoms in the bilinear attention map, suggesting these atoms represent structural features critical for ligand-protein binding. In contrast, MolUNet++ integrates substructure recognition with a substructure mask explainer to analyze important structural components.

Figure 6 presents the interpretability analysis of DrugBAN and MolUNet++. For 6QL2, MolUNet++ accurately identified the interacting benzothiazole scaffold (which is involved in an aromatic-H interaction with Leu198) and the sulfonamide region. In contrast, the highlighted explanations provided by DrugBAN were less consistent with the crystal structure interaction map, erroneously attributing significance to the ethoxy group of ethoxzolamide.

**Fig. 6.**
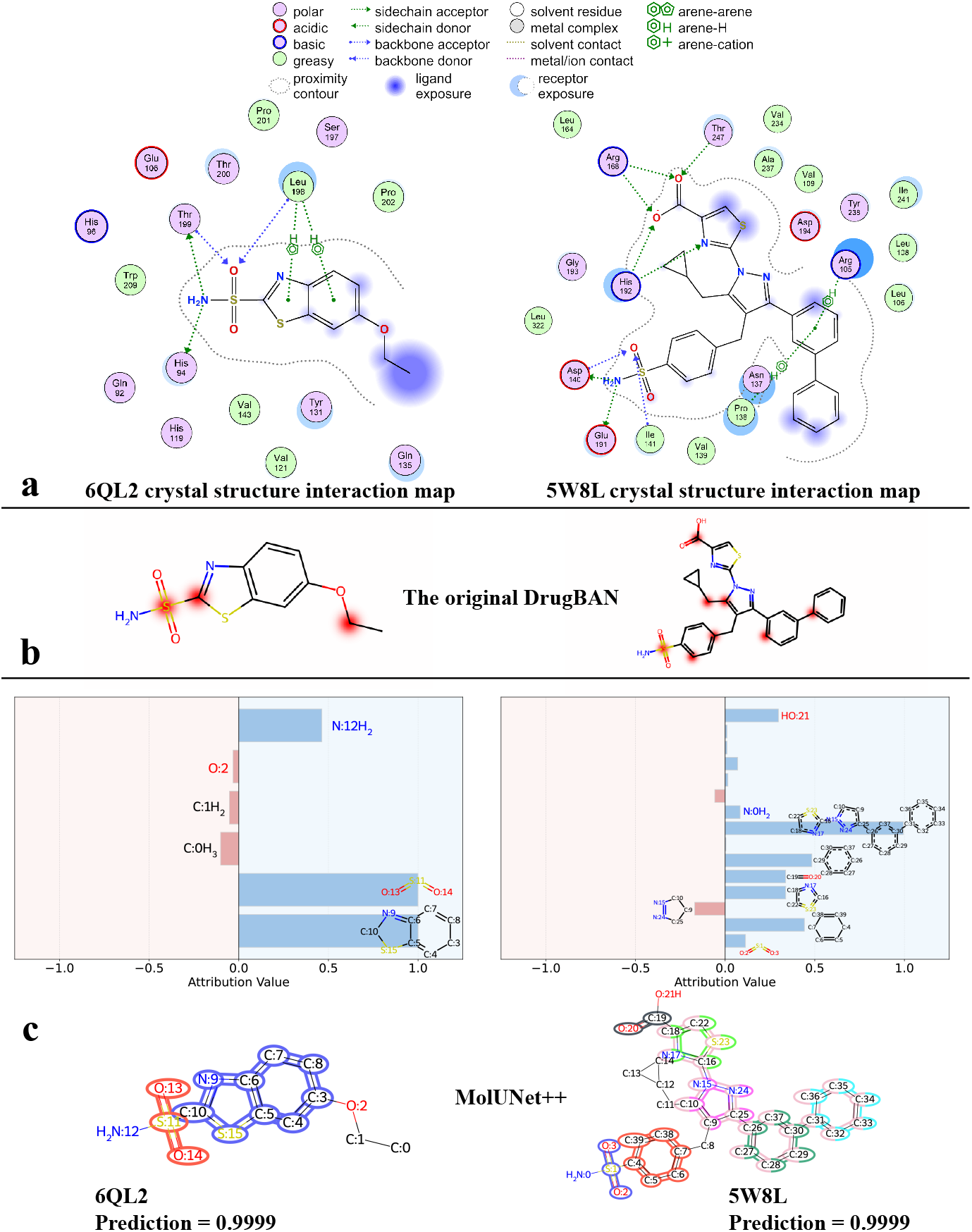
Visualization of the case study in DTI prediction. (a) The ligand-protein interaction from the corresponding crystal structures (visualized through the Molecular Operating Environment (MOE) software); (b) visualization of the most important atoms in the bilinear attention layer of DrugBAN, that contribute to the binding; (c) pooling results and identified substructure contribution visualization in MolUNet++

A similar trend can be observed in the 5W8L example. The scaffold of the ligand was assigned the highest importance by MolUNet++, while interacting fragments such as the biphenyl ring, the carboxyl group, and the sulfonamide region exhibited significant attribution values. In contrast, noninteracting structures did not display notable positive values, further supporting the accuracy of MolUNet++ in identifying critical interaction regions.

These results further confirm the effectiveness of MolUNet++ in substructure recognition, aligning with chemical logic. Its ability to identify critical interactions highlights its potential to assist in drug design by uncovering unknown interaction patterns.

## Conclusions

In this study, we propose a novel molecular representation learning model, MolUNet++, which demonstrates significant advancements over traditional GNN encoders. By utilizing graph pooling for down-sampling and up-sampling, along with a query-guiding mechanism, and employing a Nested UNet structure to integrate atomic features effectively. The framework endows MolUNet++ with the ability to recognize adaptive-grained substructures and interactions explicitly.

We evaluated the performance of MolUNet++ on three tasks: molecular property prediction, DDI prediction, and DTI prediction. Across all tasks, MolUNet++ demonstrated significant performance improvements compared to baseline models. Additionally, we conducted several case studies to analyze the interpretability of MolUNet++, where the integration of structure recognition and mask explanation offers robust chemical analysis capabilities. We believe that this interpretability can be beneficial for researchers involved in lead compound design and optimization.

This study focused on the representation learning of molecular 2D structures. We consider integrating MolUNet++ with other models based on 1D or 3D data to further enhance molecular representation as a promising avenue for future research. Additionally, this work primarily analyzed the comparative performance of MolUNet++ and traditional GNN encoders in DDI and DTI predictions, designing task-specific and more efficient models for DDI and DTI predictions based on MolUNet++ will be another objective in our future work.

In conclusion, MolUNet++ not only provides a robust framework for molecular representation learning but also presents opportunities to enhance interpretability and adaptability in molecular analysis, with potential applications in drug discovery and chemical biology.

## Data and code availability

All the datasets used in this work are from public resources. Molecular property prediction datasets can be found at https://moleculenet.org/datasets-1; DrugBank dataset in DDI prediction can be found at https://go.drugbank.com/releases; for the DTI prediction, BindingDB dataset can be found at https://www.bindingdb.org/bind/index.jsp, BioSNAP dataset can be found at https://github.com/kexinhuang12345/MolTrans/tree/master/dataset/BIOSNAP, Human dataset can be found at https://github.com/lifanchen-simm/transformerCPI. The code and the processed datasets of this work is open-sourced and available at https://github.com/xfd997700/NestedMolUNet.

## Footnotes

1 RDKit: Open-source cheminformatics. https://www.rdkit.org/

2 PyTorch: https://pytorch.org/

3 PyG (PyTorch Geometric).: https://pyg.org/

4 MoleculeNet by the DeepChem library. https://moleculenet.org/datasets-1

5 DrugBank dataset: https://go.drugbank.com/releases

